# Rapid Prototyping of Cell Culture Microdevices Using Parylene-Coated 3D Prints

**DOI:** 10.1101/2021.08.02.454773

**Authors:** Brian J. O’Grady, Michael D. Geuy, Hyosung Kim, Kylie M. Balotin, Everett R. Allchin, David C. Florian, Neelansh N. Bute, Taylor E. Scott, Gregory B. Lowen, Colin M. Fricker, Scott A. Guelcher, John P. Wikswo, Leon M. Bellan, Ethan S. Lippmann

## Abstract

Fabrication of microfluidic devices by photolithography generally requires specialized training and access to a cleanroom. As an alternative, 3D printing enables cost-effective fabrication of microdevices with complex features that would be suitable for many biomedical applications. However, commonly used resins are cytotoxic and unsuitable for devices involving cells. Furthermore, 3D prints are generally refractory to elastomer polymerization such that they cannot be used as master molds for fabricating devices from polymers (e.g. polydimethylsiloxane, or PDMS). Different post-print treatment strategies, such as heat curing, ultraviolet light exposure, and coating with silanes, have been explored to overcome these obstacles, but none have proven universally effective. Here, we show that deposition of a thin layer of parylene, a polymer commonly used for medical device applications, renders 3D prints biocompatible and allows them to be used as master molds for elastomeric device fabrication. When placed in culture dishes containing human neurons, regardless of resin type, uncoated 3D prints leached toxic material to yield complete cell death within 48 hours, whereas cells exhibited uniform viability and healthy morphology out to 21 days if the prints were coated with parylene. Diverse PDMS devices of different shapes and sizes were easily casted from parylene-coated 3D printed molds without any visible defects. As a proof-of-concept, we rapid prototyped and tested different types of PDMS devices, including triple chamber perfusion chips, droplet generators, and microwells. Overall, we suggest that the simplicity and reproducibility of this technique will make it attractive for fabricating traditional microdevices and rapid prototyping new designs. In particular, by minimizing user intervention on the fabrication and post-print treatment steps, our strategy could help make microfluidics more accessible to the biomedical research community.

## Introduction

Cell culture microdevices are traditionally designed and prototyped using photolithographical techniques performed in a cleanroom^1^. Here, photoresist is first spin coated onto a silicon wafer at a desired thickness, followed by exposure of ultraviolet light through a mask that contains patterned features. Uncured photoresist is then washed away, leaving features at a height dictated by the thickness of the original thin film. The iterative process of photoresist layering and selective curing through a series of masks ultimately builds the master mold with multilayered, planar features. To create the final microdevice, polydimethylsiloxane (PDMS) or other precursors are polymerized on the master mold, which generates the microfeatures on the resulting elastomer. Such microdevices have been workhorses for biological experiments, including those involving morphogen gradients, migration, and clonal analyses^2^. However, the technical prowess needed to fabricate master molds (particularly ones with multiple masks that need to be precisely aligned), as well as the necessary access to a cleanroom and specialty training, has limited widespread adoption of microdevices by the broad biological community despite lofty goals of open-source sharing of device designs^3^.

3D printing is a promising strategy to overcome these challenges because the fabrication process is automated (requiring only a design file) and can generate complex features in all planes. Furthermore, even entry level 3D printers can achieve micron-sized features that would be suitable for many of the applications listed above. However, most resins used for 3D printing are cytotoxic due to the inability to remove all uncured monomers and photoinitiator compounds from the print, which limits their use in direct print microdevices. In addition, uncured monomers and sequestration of catalysts can inhibit elastomer crosslinking, which prevents high-fidelity replication of intended structures.^4–6^ Various post-print treatments have been tested, but none have universally solved these issues^1,7–10^. For example, in a recently published manuscript, 16 commercially available resins were printed and subjected to various treatments before use in PDMS casting^10^. It was found that most resins, when baked and treated with ultraviolet light, could produce PDMS casts that retained high-fidelity features. However, this strategy required high temperatures and long light exposures, which can warp 3D prints and render them brittle, and the prints were not tested for biocompatibility. Another approach to post-print treatment of devices is surface coatings. In one example, 3D prints were air brushed with a protective ink to create a physical barrier, but the ink was manually applied and, as stated by the authors, practice was necessary for creating an optimum finish^11^. In other examples, 3D prints were coated with silane to create a hydrophobic fluorinated barrier on the print, which allows PDMS to be demolded^5^,^6^. Yet, multiple processing steps were still required for this strategy, and the toxicity of silane also prohibits prints from being used directly for biological applications. Overall, while progress has been made towards making 3D prints biocompatible and suitable as master molds^4–13^, more effective and easily implemented methods are still needed.

As a potential solution to this problem, we explored the use of parylene coatings on 3D prints. Parylene is an FDA USP Class VI polymer that meets the strictest testing requirements for human implantation. Nanometer-thick films are commonly deposited on implanted medical devices to create a hydrophobic barrier that is impermeable to small molecules, water, and gases; this hydrophobic barrier preserves the long-term integrity of devices in the human body^12,13^. Parylene films have also been used to prevent small molecule permeation through PDMS microdevices in biosensing applications^14,15^. Hence, we hypothesized that parylene deposition on high-resolution stereolithography (SLA) 3D prints could serve two purposes: parylene would render SLA 3D prints cytocompatible and allow SLA 3D prints to serve as master molds for PDMS casting. Here, we show these hypotheses hold true and demonstrate the utility of parylene-coated 3D prints for a wide variety of applications.

## Methods

### Microdevice fabrication

All prints were designed with Fusion 360 (Autodesk) and fabricated using a Form 3 SLA printer (Formlabs). Fusion 360 designs were converted to STL files. The slicing and coding of the STL file was converted into g-code using PreForm software (Formlabs). The Form 3 printer provides a minimum layer resolution of 25 μm along the z-axis and 80 μm resolution along the x- and y-axes. Four different resins were used in this manuscript: Grey, Clear, Black, and High Temp (Formlabs). Completed prints were washed with isopropyl alcohol until visibly clean, then placed in the Formlabs UV cure for 60 seconds at 60°C. Cured prints were then transferred to a Labcoter PDS 2010 parylene deposition machine (Specialty Coating Systems) and coated with parylene-C according to the manufacturer’s instructions. If being used directly for cell culture, prints were sterilized using Gamma irradiation with dose rays of 2.5 × 10^−3^ J g^−1^ s^−1^ for 12 hours. Otherwise, prints were placed in a plastic petri dish, and polydimethyl siloxane (PDMS, Sylgard 184; Ellsworth Adhesive Company) elastomer and curing agent were mixed at a weight ratio of 10:1 and poured onto the print. The mixture was then degassed for 30 minutes in a vacuum chamber and cured in an oven at 80°C for 3 hours. After demolding, PDMS devices were bonded to glass coverslips using a PlasmaFlo PDC-FMG plasma cleaner (Harrick Plasma) and sterilized in an autoclave prior to use.

### Scanning electron microscopy

Rectangular prints with 2 mm length were designed and printed. Lab tape was placed over one half of the print, followed by coating parylene-C. The tape was then removed, and the prints were mounted on a Ted Pella pin mount. To characterize the interface of deposited parylene-C and exposed print, the samples were imaged using a scanning electron microscope (Zeiss Merlin) at an accelerating voltage of 2 kV. The thickness of the parylene layer was measured using the Zeiss software.

### Curvature measurements

Angular wall features, specifically the width of angular channels on a 3D print and its corresponding PDMS device, were measured using a Veeco Daktak 150 profilometer (Bruker).

### Cell maintenance

CC3 human induced pluripotent stem cells (iPSCs) were maintained in E8 medium on standard tissue culture plates coated with growth-factor reduced Matrigel (Corning). iPSCs were passaged at 60-70% confluence using Versene (Thermo Fisher Scientific). Jurkat E6-1 cells were cultured in suspension in RMPI 1640 medium containing 2 mM glutamine, 10% fetal bovine serum (FBS), 100 U mL ^−1^ penicillin, and 50 μg mL^−1^ streptomycin. MDA-MB-231 cells were cultured in DMEM supplemented with 10% FBS, 100 U mL^−1^ penicillin, and 50 μg mL^−1^ streptomycin. Primary human mesenchymal stem cells (MSCs) were purchased from Extem Biosciences and cultured in MSC Growth Medium 2 (PromoCell). Peripheral blood mononuclear cells (PBMCs) were isolated from primary donors using Ficoll-Paque Plus (GE Life Sciences) and frozen until use. All cells were maintained at 37°C in a humidified atmosphere of 5% CO_2_.

### Direct print neuronal toxicity assays

Cortical glutamatergic neurons were generated from iPSCs as previously described^16^. Neurons were detached from plates by a 5-minute incubation with Accutase (Thermo Fisher Scientific), centrifuged, and pipetted into a Matrigel-coated 96-well plate at a density of 2×10^5^ cells mL^−1^. After allowing the neurons to recover for 2 weeks with media changes every 2-3 days, uncoated and parylene-coated prints were then immersed in each well. At 1, 2, 7, and 14 days after immersion, separate wells of neurons were incubated with 1 μM CytoCalcein Violet 450 (AAT Bioquest) and 1 μM propidium iodide (Thermo Fisher Scientific) for 1 hour, followed by imaging on a Zeiss LSM 710 confocal microscope and quantification with ImageJ. After 21 days, the remaining neurons were fixed in 4% paraformaldehyde for 10 minutes and washed with PBS. A solution of 5% goat serum and 0.03% Triton X-100 was used to block and permeabilize the cells overnight on a rocking platform at room temperature. Cells were washed again in PBS and incubated overnight with 4’,6-diamidino-2-phenylindole (DAPI; Thermo Fisher Scientific) and Alexa Fluor 647-conjugated βIII tubulin (Abcam ab190575). After final PBS washes, cells were imaged on a Zeiss LSM 710 confocal microscope.

### Bone on chip perfusion device

A nanocrystalline hydroxyapatite (nHA)-poly(ester urethane) (PEUR) composite foam scaffold, fabricated as previously described^17^, was cut to a size of 5 mm by 5 mm by 2 mm and wedged into a perfusion device consisting of a single channel with dimensions of 50 mm by 5 mm by 2 mm. Human MSCs, PBMCs, and MDA-MB-231 cells were labeled with fluorescent CellTracker™ (Thermo Fisher Scientific) membrane dyes (blue, green, and deep red, respectively) and seeded within the scaffold. The scaffold was cultured for 7 days under static conditions in α-MEM media containing 50 μg mL^−1^ L-ascorbic acid, 10 nM dexamethasone, 10 mM β-glycerophosphate, 10 nM Vitamin D, 25 ng mL^−1^ macrophage colony-stimulating factor, and 50 ng mL^−1^ receptor activator of nuclear factor kappa-B ligand (Sigma Aldrich). The scaffold was then perfused at 56 μL per second for 28 days in the same medium. The fluorescently-labeled triculture was imaged periodically with a Zeiss LSM 780 confocal microscope for 14 days after which the CellTracker™ dyes were no longer visible. At defined time points, 1 μM of PerkinElmer® OsteoSense® fluorescently labeled bisphosphonate was added to the perfused media to visualize mineralization.

### Droplet assays

As described in the main text, 3D printed microfluidic devices were designed based on previous publications,^18^ with a trial-and-error approach until the final designs yielded desired properties. Biopsy punches were used to create the inlets and outlet from the PDMS devices after plasma bonding to a glass slide. Solutions were loaded into glass Hamilton syringes and pumped through devices with Harvard Apparatus Pump 11 Pico Plus Elite syringe pumps. 10 mL of aqueous solution and 10 mL of oil solution were used for all experiments. For the single-wall emulsion experiments, the aqueous phase consisted of 10 μM fluorescein in PBS and the oil phase was pure mineral oil. Flow rates for each phase were adjusted as indicated in the figure legend. Droplets were imaged using a Zeiss Axiovert 200 microscope and diameter was quantified using ImageJ after calibration with a stage micrometer slide. For the double-wall emulsion experiments involving Jurkat encapsulation, the aqueous phase consisted of PBS with 1 μM calcein and the oil phase consisted of 2.2% Ionic Krytox 157 FS-H in HFE7500. Cell density was 3.6×10^6^ cells per mL and flow rates were 50 μL per minute for the aqueous solution and 100 μL per minute for the oil solution. For the double-wall emulsion experiments involving aptamer encapsulation, the oil phase consisted of 2.2% Ionic Krytox 157 FS-H in HFE7500 and the aqueous phase consisted of 40 μM DFHBI (Sigma-Aldrich) with or without 50 nM Broccoli aptamer in binding buffer (40 mM Tris HCl, 280 mM NaCl, 10 mM KCl, 1 mM MgCl2, and 2 mM CaCl2 in water). Flow rates were 50 μL per minute for the aqueous solution and 100 μL per minute for the oil solution. Broccoli aptamer was produced starting with plasmid pAV-U6+27-Tornado-Broccoli (gifted by Dr. Samie Jaffrey; Addgene plasmid 124360). PCR was performed using Phusion high-fidelity polymerase (NEB) to add a T7 promoter sequence with custom primers (forward: GTATAATACGACTCACTATAGGGAAACCGCCTAACCATGCCG; reverse: GGCATTGGCAGTGTTCTACAGTCC). PCR fragments were then *in vitro* transcribed using the AmpliScribe T7-Flash kit (Lucigen) with overnight incubation at 37°C, and the resultant RNA was ethanol precipitated overnight at −20°C before resuspension in nuclease-free water and storage at −20°C. Prior to experiments, RNA was thawed and diluted to the aforementioned concentration in binding buffer. RNA in binding buffer was heat pre-treated at 98°C for 15 minutes and then cooled to room temperature to ensure proper aptamer folding. RNA was then incubated with DFHBI for 15 minutes before use in the droplet generator.

### Brain organoid assays

Aggrewell™ 800 (Stemcell Technologies) or size-matched PDMS microwell arrays were placed in 12-well plates. CC3 iPSCs were collected using Accutase and seeded at a density of 3×10^6^ cells per well. Plates were centrifuged at 100xg for 3 minutes in E8 medium containing 10 μM Y27632 (Tocris). The following day, the medium was switched to E6 medium supplemented with 10 μM SB431542 (Tocris) and 0.4 μM LDN1931189 (Tocris) for 6 days to induce neuralization. Neural organoids were then removed from the PDMS microwell arrays were placed in a petri dish and imaged using an EVOS brightfield microscope. Circularity was quantified using ImageJ.

For fusion experiments, on day 7, the organoids were transferred from the aforementioned PDMS microwells to a previously published spinning bioreactor^19^ with a speed of 80 revolutions per minute in E6 medium containing 10 ng mL^−1^ FGF2 (Peprotech), 20 ng mL^−1^ BDNF (Peprotech), and 20 ng mL^−1^ GDNF (Peprotech) with or without activation of sonic hedgehog signaling via 1 μM purmorphamine (Cayman Chemical). This approach mimics a previous publication generating dorsal and ventral forebrain organoids^20^. At day 14, ventral organoids were incubated with 200 nM Tubulin Tracker™ Deep Red (Thermo Fisher Scientific) for 1 hour at 37°C with 5% CO_2_. The organoids were washed 3 times by PBS before seeding into the “fusion” PDMS microwell array housed in a 12-well plate. Approximately 120 organoids were manually added to the array at a 1:1 ratio of dorsal and ventral organoids and allowed to settle in the microwells. Organoids were incubated in the fusion microwells for 2 days in E6 medium containing FGF2, BDNF, and GDNF, and then transferred to the spinning bioreactor in the same medium for an additional 2 days. Afterwards, organoids were divided into smaller groups and imaged using a Zeiss LSM 710 confocal microscope. The organoids were manually counted to quantify distributions of fusion events.

## Results

In a standard commercial parylene coater, parylene dimers are sublimated, converted to monomers through pyrolysis, and then deposited on surfaces for polymerization (**Figure 1A**). Using scanning electron microscopy, we verified that we could deposit parylene on the surface of prints fabricated with an entry level SLA 3D printer (**Figure 1B**). We then crosslinked PDMS on uncoated versus parylene-coated prints and observed facile demolding solely from the coated prints, leaving smooth, curved, micropatterned features (**Figure 1C**). We further used profilometry to show that curvature on the 3D print and resultant PDMS device were perfectly aligned (**Figure 1D**). This was especially notable because curved, non-planar features are extremely difficult to achieve by photolithography. To highlight the rapid prototyping capabilities of this approach, we generated variations of a triple-chamber microdevice with different heights and pillar spacing^21^. All devices were designed with computer-aided design (CAD) Fusion360 software, 3D printed, and cast with PDMS in a single day (**Figure 2**), and the final device reproducibly achieved 180 μm spacing between adjacent pillars without any noticeable defects.

**Figure 1.**
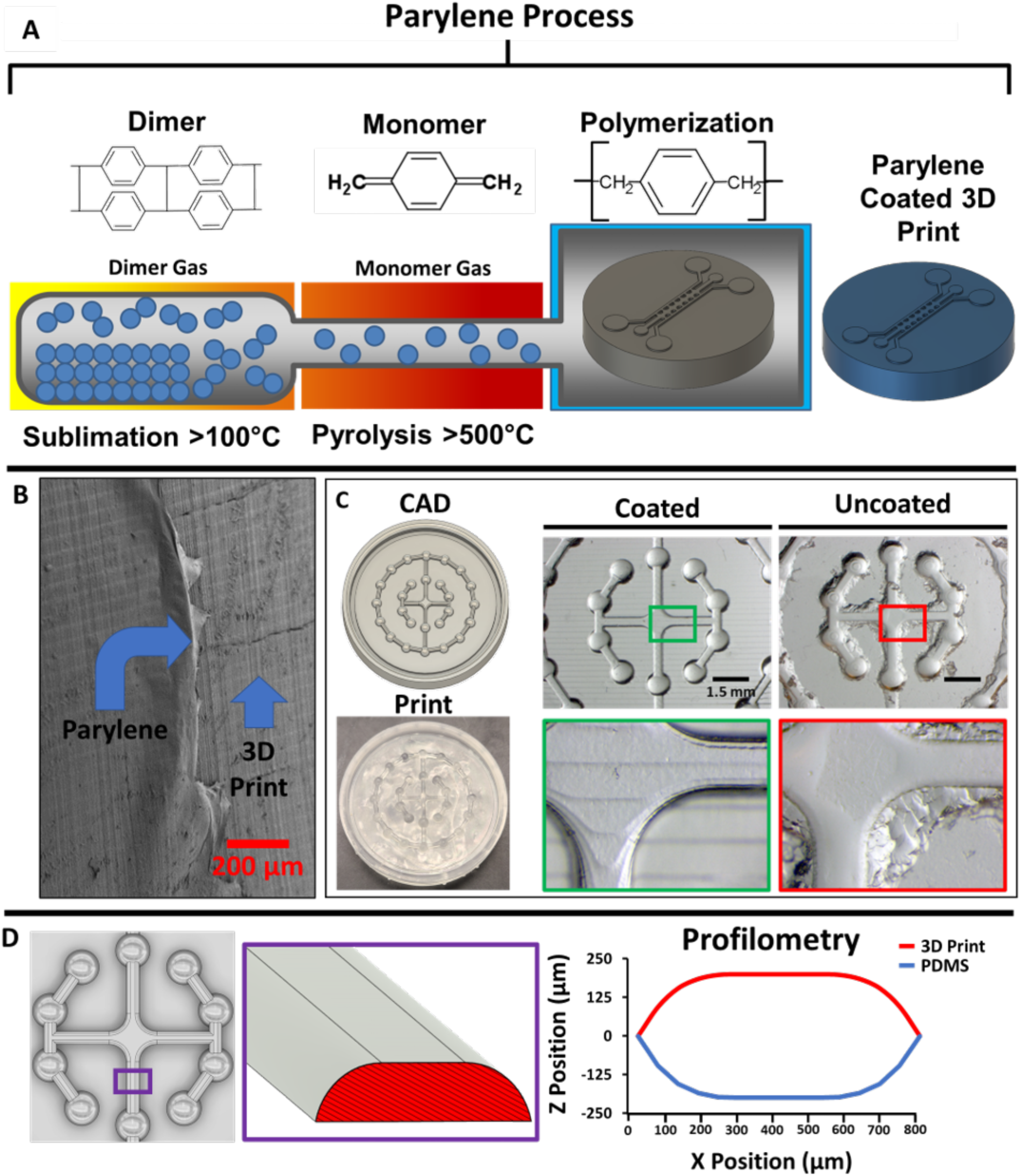
Parylene deposition renders 3D prints amenable to soft lithography. **A)** Overview of the parylene vapor deposition process. **B**) Scanning electron microscopy image of a 3D print coated with an 18 nm thick layer of parylene. **C**) Example of PDMS demolding from a 3D print with complex features. Cast PDMS from the parylene-coated 3D print demonstrated facile demolding, high-fidelity contouring of the non-planar features, and fully crosslinked PDMS. Cast PDMS from the uncoated prints was difficult to demold and retained minimal contouring of the non-planar features with evidence of residual material that was not fully crosslinked. **D**) Measured high angle profile of a select portion (purple box) of the angular walls on the 3D print and casted PDMS mold.

**Figure 2.**
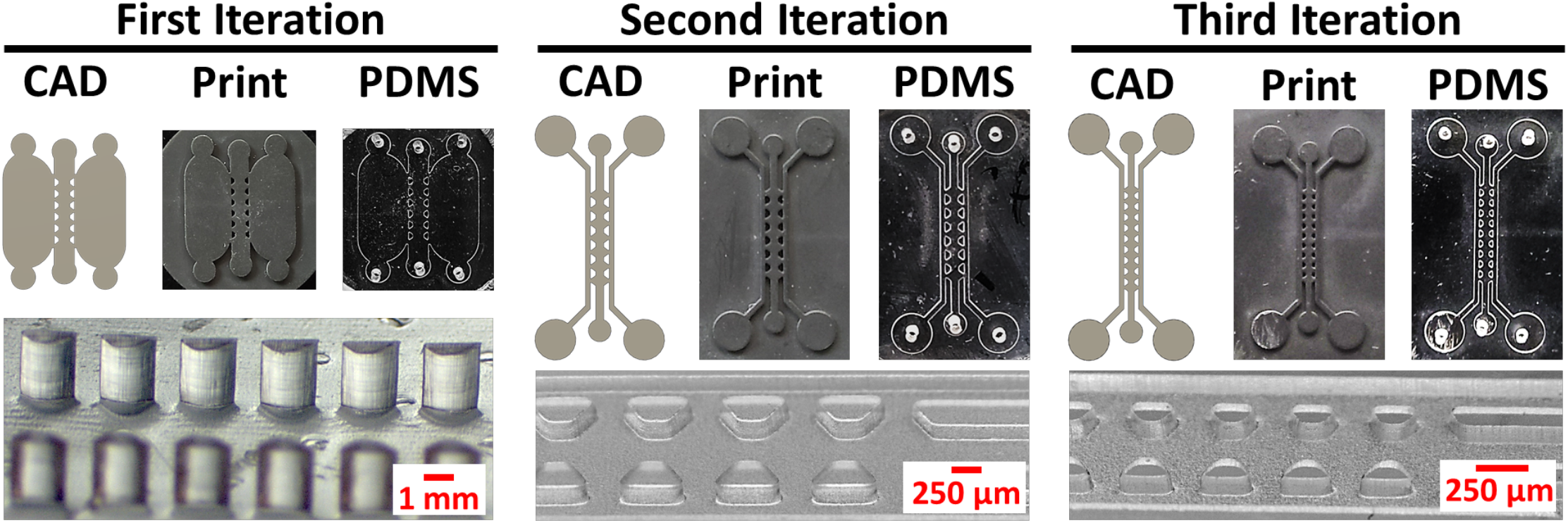
Rapid prototyping of triple chamber microdevices from parylene-coated 3D prints. Each iteration shows the CAD file, 3D print, and the final casted PDMS device. Brightfield images of pillar height, shape, and spacing are shown to highlight differences in the microfeatures.

Having shown the prototyping capabilities of this method, we verified its applicability to cell culture. Using multiple different SLA resins, we 3D printed a series of pillars that could interface with a 96-well plate containing human induced pluripotent stem cell (iPSC)-derived excitatory neurons (**Figure 3A**). Submersion of uncoated pillars in media resulted in widespread neuron death after 2 days, presumably due to monomer leaching from the prints (**Figure 3B-C**). In contrast, independent of resin type, parylene coating rendered the prints cytocompatible with >95% neuron viability after 2 weeks (**Figure 3B-C**). Of note, neurons in wells with parylene-coated prints exhibited healthy morphology with robust extension of βIII-tubulin+ neurites after 21 days (**Figure 3D**). To showcase this approach for simple applications, we fabricated a custom millifluidic channel to fit a millimeter-sized polymer construct as a trabecular-bone-on-a-chip model. This device was not commercially available, and a vendor quoted fabrication at ~$15,000 for a minimum order of 20 custom micromachined devices and a lead time of 6 months. In contrast, we fabricated the device in-house in less than 6 hours with material costs of less than a dollar. The construct was perfused within the flow channel, yielding long-term cell survival and mineralization of the construct after several weeks (**Supplementary Fig. 1**).

**Figure 3.**
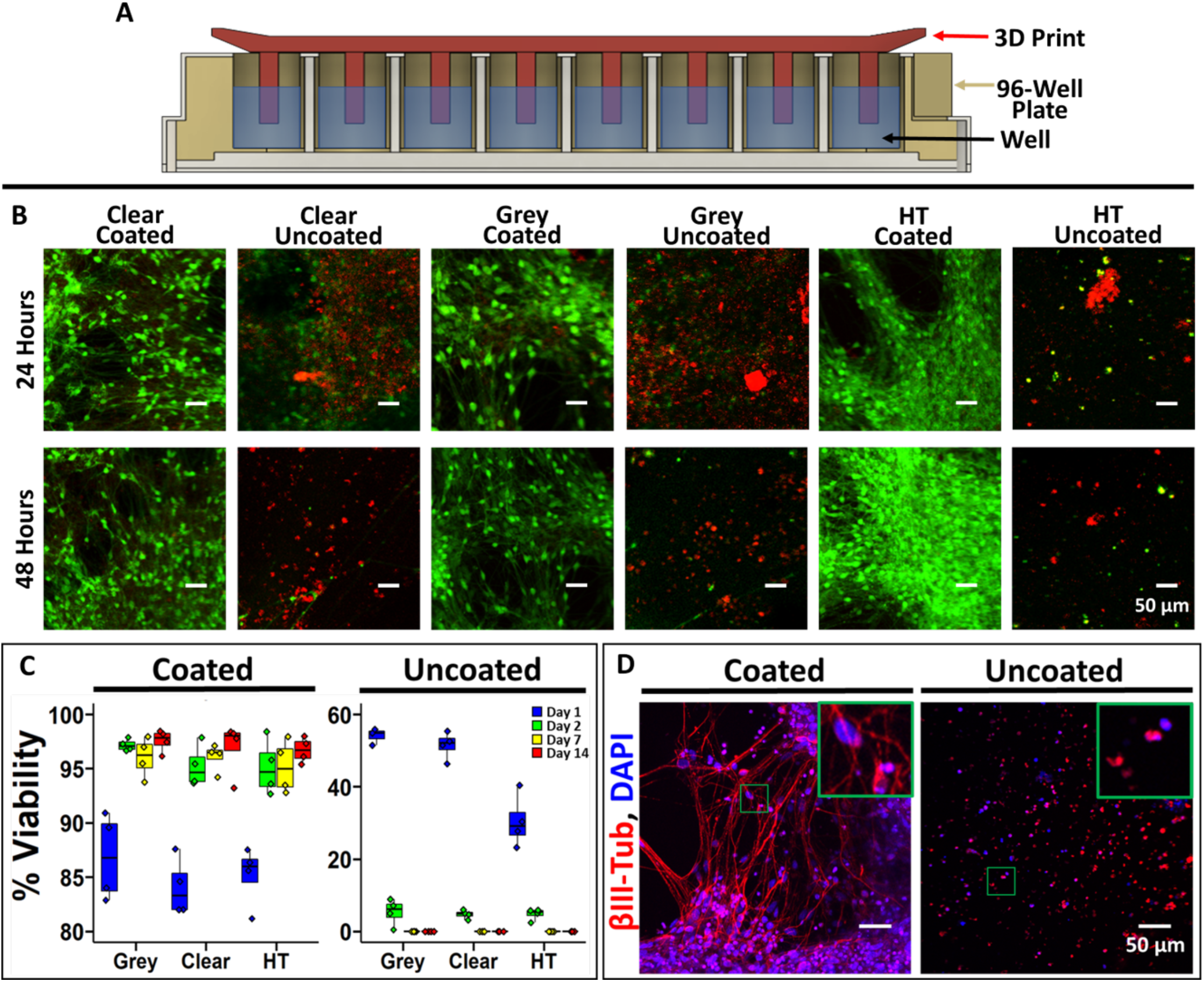
Biocompatibility of parylene-coated 3D prints. **A)** Schematic cross-sectional representation of a 96-well plate with the 3D printed insert used for biocompatibility testing. **B)** Representative images of iPSC-derived neurons labeled with Calcein (live cells) and propidium iodide (dead cells) after 24 and 48 hours of soluble contact with the 3D prints. **C)** Quantification of iPSC-derived neuron viability at different time points (percent of live cells versus total cells). Each well was imaged in 3 different locations, and 4 independent wells were imaged per condition representing the individual data points. On day 14, a one-way ANOVA was used to calculate statistical significance between the coated and uncoated conditions (p<0.0001). **D)** Representative immunofluorescence images of the iPSC-derived neurons at day 21 after incubation with coated and uncoated 3D prints (grey resin). Inset highlights morphological differences.

To further show the utility of our approach, we next applied it towards microfluidic droplet generators, which are frequently used to encapsulate biological materials for high throughput investigations^18^. We first prototyped a single-wall emulsion, flow-focusing PDMS droplet generator casted from a 3D printed, parylene-coated mold (**Figure 4A-B**). We demonstrated tight control over droplet size by altering the flow rates (**Figure 4C**). We next prototyped a double-emulsion droplet generator with slightly more complicated features. From a single 3D printed, parylene-coated flow-focusing mold (**Figure 4D**), we developed a PDMS droplet generator capable of producing double-wall emulsions, which we subsequently used to encapsulate Jurkat cells (**Figure 4E**). These cells were fluorescently labeled with the live-cell marker Calcein-AM, permitting easy identification within the innermost phase of the bi-layered droplets. As an additional proof-of-concept, we encapsulated the chromophore 3,5-difluoro-4-hydroxybenzylidene imidazolinone (DFHBI) with its cognate RNA aptamer Broccoli^22^, showing increased fluorescence of the chromophore when the aptamer is present in the droplets (**Figure 4E-F**).

**Figure 4.**
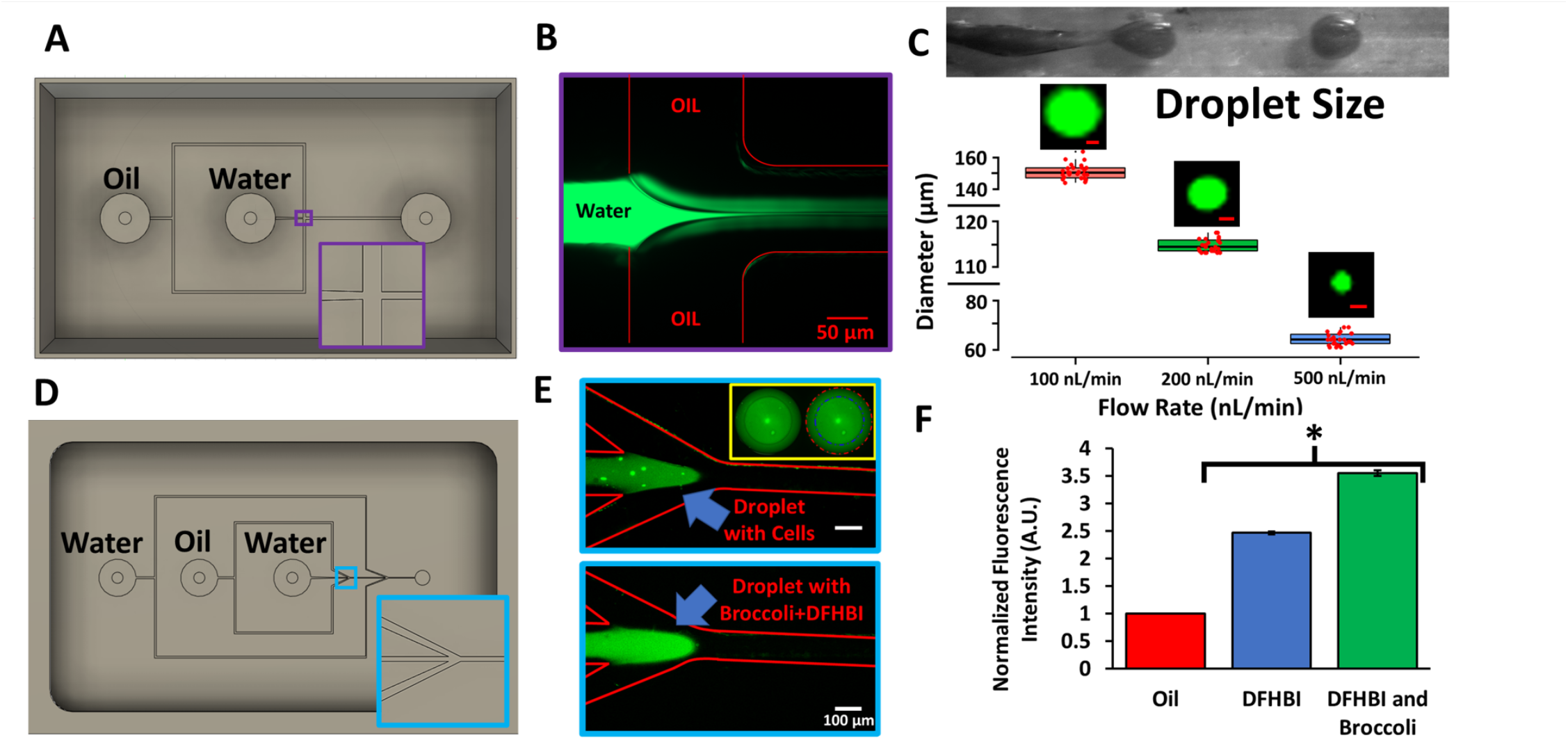
PDMS droplet generators cast from 3D printed molds. **A**) CAD design for the 3D printed single-wall emulsion droplet generator. Inset highlights the geometry of the flow-focusing T-junction. **B**) Representative image of the fluorescein-filled aqueous phase jetting into the oil phase to create the emulsion. **C**) Brightfield image of the jetting regime and subsequent droplet generation. At 3 different flow rates, droplets diameters were quantified using ImageJ. Data are presented as mean and standard deviation from 3 independent replicates (~100 droplets were counted per replicate, and randomly selected droplets are shown as individual red data points to highlight reproducibility). Representative fluorescein-filled droplets are shown for each flow rate (scale bars, 50 μm). **D**) CAD design for the 3D printed double-wall emulsion droplet generator. Inset highlights the angled flow-focusing junction. **E**) Representative images of the jetting regime where the aqueous phase contains Calcein-labeled Jurkat cells or the Broccoli RNA aptamer with DFHBI. The inset highlights the double-wall shell containing labeled Jurkat cells (blue line, inner wall; red line, outer wall). **F**) Quantification of fluorescence of double-wall droplets containing DFBH versus DFHB and the Broccoli RNA aptamer. Fluorescence was normalized to the oil phase. Data are presented as mean and standard deviation from 3 independent replicates. A student’s unpaired t-test was used to compare DFHBI with and without Broccoli (*p<0.0001).

Last, to demonstrate the utility of this technique for rapid prototyping of novel, more complex devices, we fabricated custom cell culture microwells. We designed 3D printed pyramids to mimic the dimensions of commercially available plastic microwell inserts (i.e. Aggrewell™ 800) (**Figure 5A**). PDMS microwells cast from these pyramids were seeded with iPSCs in culture media that directs differentiation to brain organoids. The iPSCs readily aggregated in the PDMS microwells and there was no difference in shape compared to brain organoids produced in commercial Aggrewells™ (**Figure 5A**). Because Aggrewells™ are only available in two sizes of the same design, we highlighted the iterative capability of our approach by designing a narrower, deeper microwell structure to enable high throughput generation of “assembloids” (fused organoids). Assembloids are normally generated by manual placement and extended contact of two organoids, which is an inherently low throughput technique^20^. We designed oblong, tapering wells to fit 2 brain organoids within a single well, which allowed us to generate large batches of dorsal/ventral forebrain assembloids in a single seeding, with a high rate of success and a uniform distribution of fusion events (**Figure 5B**).

**Figure 5.**
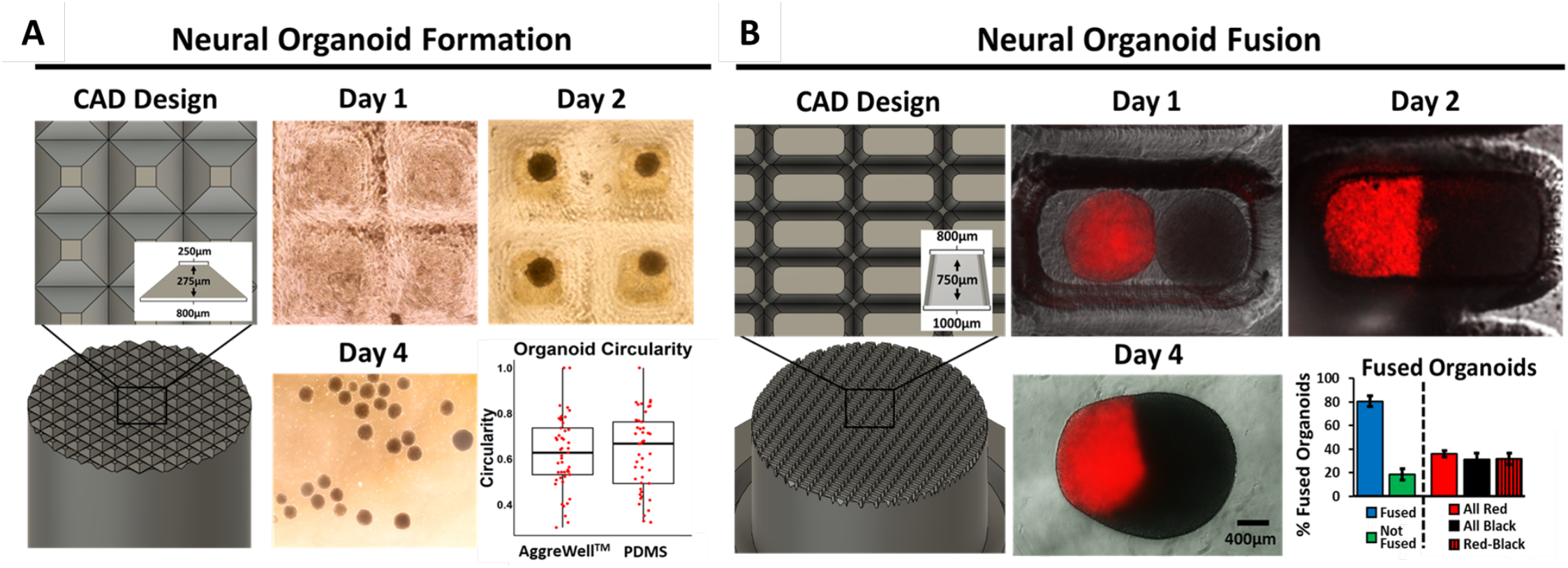
Custom 3D printed microwell arrays for batch preparations of iPSC-derived brain organoids and assembloids. **A)** 3D prints were designed with pyramids matching Aggrewell™ 800 dimensions and casted with PDMS to create microwells. Representative brightfield images are shown for day 1 (before aggregation), day 2 (after aggregation), and day 4 (after removal from the wells). Organoid circularity was quantified on day 7 and compared to organoids prepared in an Aggrewell™ 800. Each data point represents a single organoid, and measurements were pooled from 3 biological replicates. Statistical significance was calculated using the student’s unpaired t-test (p>0.05). **B)** Based on the sizes of organoids in panel a, 3D prints were designed to create wells that would hold 2 organoids (1:1 ratio of unlabeled dorsal forebrain organoids and Tubulin Tracker™ Deep Red-labeled ventral forebrain organoids). Representative fluorescence microscopy images are shown on day 1, day 2, and day 4 after organoid seeding to highlight fusion events. The number of fused versus unfused organoids, as well as the distribution of fused organoids, was quantified on day 10 after fusion. Data are presented as mean and standard deviation from 3 pooled biological replicates (98 total events counted).

## Discussion

Overall, we present a simple, cost-effective strategy for rapid prototyping cell culture microdevices that eliminates the need for cleanroom access and specialized training. To fit the specific needs of researchers seeking to understand subtle nuances of biology, microdevices need to become more modular and personalized. Yet, most commercial products are generated in a one-size-fits-all format without options for customization. Our approach provides universal freedom to iteratively adapt devices to a specific experiment without extensive cost or time investments. Furthermore, while many smaller academic institutions and companies simply cannot afford to build and maintain their own cleanroom, it is likely that some can afford a standard SLA 3D printer (~$3,000-4,000) and parylene coater (~$35,000-50,000) as part of a core facility.

Even when researchers have access to a cleanroom, the intellectual barrier for use by non-experts remains high. While CAD files for single layer and multilayer master molds can be readily shared on open-source forums, the end user must still have sufficient technical prowess to produce the mold within a cleanroom. Using our strategy, CAD files can be directly converted into a master mold on a 3D printer by users with minimal manufacturing skill. This advancement is particularly powerful for users that want to embrace iterative design and testing, since printing a mold and coating it with parylene, followed by casting a PDMS device from the mold, can be accomplished in a standard workday with minimal hands-on intervention, in stark contrast to the intricate processes of photolithography and post-print processing that require user optimization. In comparison to other strategies available for treatment of 3D prints to render them suitable for cell culture or elastomeric device casting, we believe automated parylene deposition is much more straightforward and easier to replicate than UV curing and heat treatment, which can vary depending on the resin composition of the print. In comparison to silanization approaches, parylene coating is much safer, easier to handle, self-contained within the machine, and does not require a heating step that can warp and fracture the 3D prints. Given their application in implanted medical devices, parylene coatings are also expected to be much more stable than techniques that coat with inks. Anecdotally, we have used a single parylene-coated 3D print continuously over a 12-month period without any loss of fidelity in the PDMS casts.

Ultimately, we suggest that this simple and universal method has the potential to “democratize” cell culture microdevice fabrication and the use of microfluidics for biomedical research. This strategy could also broadly enable research, educational, and commercial initiatives, and prove particularly empowering for users that lack an engineering background and at institutions with minimal resources. Furthermore, while we could only achieve features with resolution of ~150 μm using our entry-level 3D printer, we note that the resolution of 3D printers continues to improve (for example, 2-photon printers that achieve nanometer features^23^). Hence, in the future, we speculate that this strategy could complement photolithography for a diverse array of applications that require single micron resolution.

## Author contributions

BJO conceived the method, designed all microdevices, and conducted data analyses. MDG performed crucial proof-of-concept experiments. HK, KMB, ERA, DCF, NNB, TES, GBL, and CMF assisted with biological experiments and data analyses. SAG provided experimental input. JPW and LMB provided access to key equipment. ESL helped design most of the experiments and supervised the research. BJO and ESL wrote the majority of the manuscript, with all authors providing feedback and revision.

## Conflicts of interest

There are no conflicts to declare.

## Acknowledgments

Funding was provided by the Chan Zuckerberg Initiative (grant 2018-191850 to ESL), the National Institutes of Health (grants R01NS110665 to ESL, R61NS112445 to ESL, R21NS116257 to LMB, and RF1MH123971 to LMB), and the National Science Foundation (grants 1846860 and 1706155 to ESL). BJO was supported by the Vanderbilt Interdisciplinary Training Program in Alzheimer’s Disease (T32 AG058524). KMB was supported by the Training Program in Environmental Toxicology (T32 ES007028). The authors would like to thank Anna Combes for her help coding in R Studio.

## Notes

### Competing Interest Statement

The authors have declared no competing interest.

