## Supplemental information for "Rapid Prototyping of Cell Culture Microdevices Using Parylene-Coated 3D Prints"

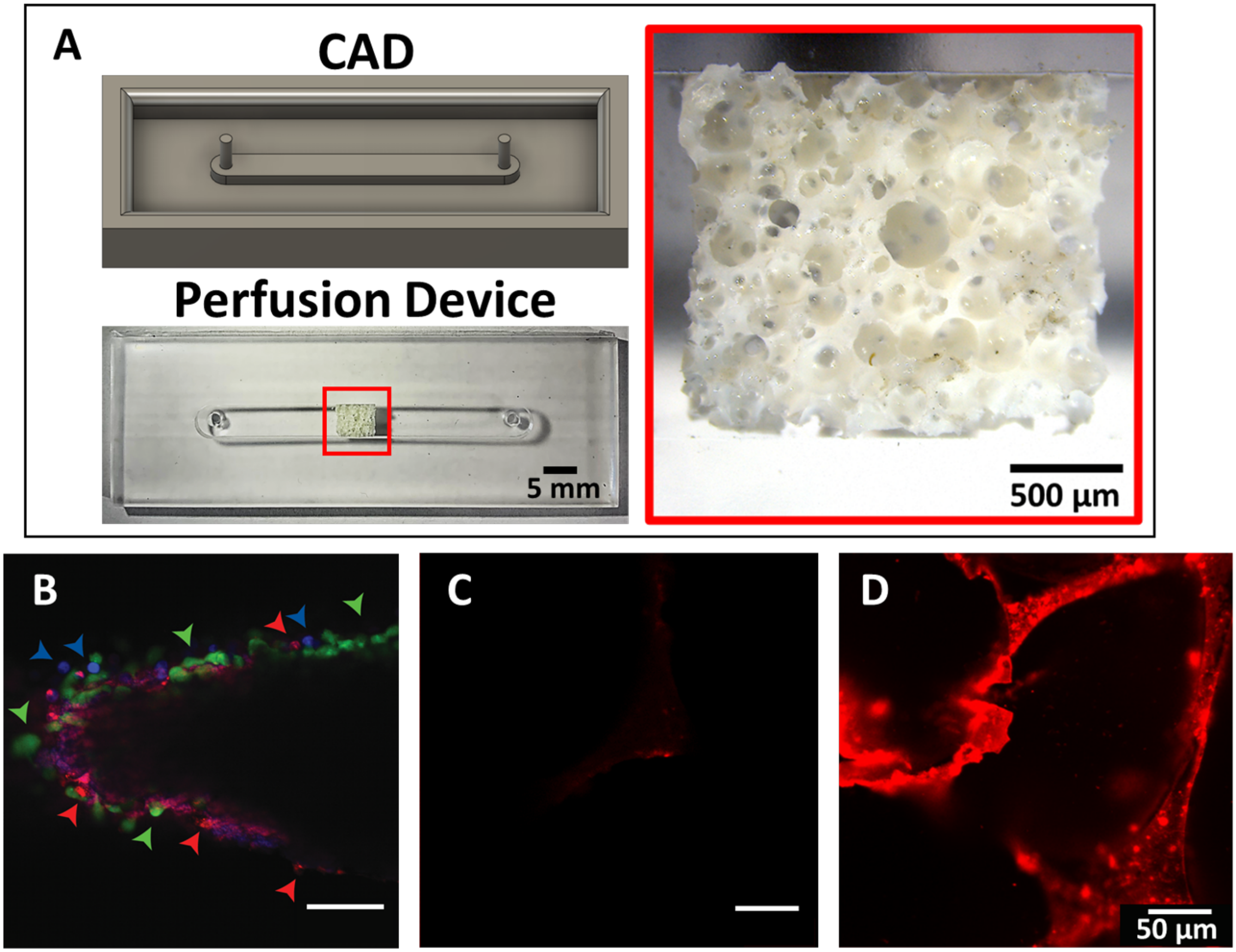


**Supplementary Figure 1**. **Trabecular bone-on-chip perfusion model. A**) A trabecular-bone-on-a-chip device was generated by fitting a noncrystaline hydroxyapatite poly(ester urethane) (nHA-PEUR) foam inside of a custom perfusion channel. **B**) Images of human MSCs (red), PBMCs (blue), and MDA-MB-231 cells (green) labeled with fluorescent CellTracker$^{TM}$ membrane dyes, 7 days after seeding in the nHA-PEUR scaffold. **C-D**) After 21 days, perfused scaffolds lacking human MSCs exhibit no mineralization (panel C) whereas scaffolds containing human MSCs exhibit robust mineralization (red, panel D).
